# Suppression of the *E*. *coli rnpA49* conditionally lethal phenotype via different compensatory mutations

**DOI:** 10.1101/2023.12.05.570125

**Authors:** Arianne M. Babina, Leif A. Kirsebom, Dan I. Andersson

## Abstract

RNase P is an essential enzyme found across all domains of life that is responsible for the 5’-end maturation of precursor tRNA transcripts. Since its discovery in the 1970s, numerous studies have sought to elucidate the mechanisms and biochemistry governing RNase P function. However, much remains unknown about the regulation of RNase P expression, the turnover and degradation of the enzyme, and the mechanisms underlying the phenotypes and complementation of specific RNase P mutations. In *Escherichia coli*, the temperature-sensitive *rnpA49* mutation in the protein subunit of RNase P has arguably been one of the most well-studied and commonly used mutations for examining the enzyme’s activity *in vivo*. Here we report for the first time naturally-occurring temperature-resistant suppressor mutations of *E. coli* strains carrying the *rnpA49* allele. We find that *rnpA49* strains can partially compensate the temperature-sensitive defect via gene amplifications of either RNase P subunit (*rnpA49* or *rnpB*) or by the acquisition of loss-of-function mutations in Lon protease or RNase R. Our results agree with previous plasmid overexpression and gene deletion complementation studies and importantly suggest the involvement of Lon protease in the degradation and/or regulatory pathway(s) of the mutant protein subunit of RNase P. This work offers novel insight into the behavior and complementation of the *rnpA49* allele *in vivo* and provides direction for follow-up studies regarding RNase P regulation and turnover.

## INTRODUCTION

RNase P is an ancient and essential endoribonuclease found in Bacteria, Archaea, and Eukaryotes that is responsible for the cleavage of 5’-end leader sequences present on premature tRNA (pre-tRNA) transcripts. In *Escherichia coli*, RNase P is a heterodimer comprised of a small protein subunit (C5, encoded by *rnpA*) and an RNA subunit (M1 RNA, encoded by *rnpB*) (Harris et al. 1998; Altman and Kirsebom 1999). The M1 RNA is the catalytic subunit of the enzyme and plays a role in substrate recognition (Guerrier-Takada et al. 1983), and the C5 protein is important for M1 RNA stability and has been implicated in electrostatic shielding, substrate binding, product release, and preventing re-binding of the product (Tallsjö and Kirsebom 1993; Buck et al. 2005; Sun and Harris 2007; Kirsebom and Trobro 2009; Lin et al. 2016). In addition to the 5’-end processing of pre-tRNAs, in *E. coli*, RNase P is essential for the separation of pre-tRNAs from polycistronic operon transcripts (Mohanty et al. 2020), and the enzyme is also involved in the processing of the 4.5S RNA, tmRNA, and mRNA transcripts (Bothwell et al. 1976; Alifano et al. 1994; Komine et al. 1994; Li et al. 2003; Li and Altman 2003).

The temperature-sensitive (ts) *rnpA49* allele in *E. coli* was first isolated from mutagenesis experiments in 1973 and is caused by an A to G transition resulting in the substitution of an arginine to a histidine at position 46 within the C5 protein (C5^A49^) (Schedl and Primakoff 1973; Apirion 1980; Kirsebom et al. 1988). This mutation reduces the solubility of the C5 protein and is hypothesized to impact the association of the C5 and M1 subunits to form the RNase P holoenzyme, resulting in reduced RNase P stability and activity (Baer et al. 1989; Li et al. 2003). Strains containing the *rnpA49* allele (designated as A49) demonstrate a slight growth defect at permissive temperatures (30-33°C), and rapidly accumulate precursor tRNAs and unprocessed polycistronic tRNA transcripts when shifted to the non-permissive temperature (42°C), eventually leading to arrest of cell growth and death (Schedl and Primakoff 1973; Li et al. 2003; Mohanty and Kushner 2007, 2008; Agrawal et al. 2014).

Previous work has shown that the *rnpA49* ts phenotype can be complemented by plasmid overexpression of either the wild-type or mutant C5 protein (Vioque et al. 1988; Jovanovic et al. 2002), the M1 RNA from *E. coli* (Jain et al. 1982; Motamedi et al. 1984; Baer et al. 1989), or with various C5 homologues from other bacterial species (Morse and Schmidt 1992; Pascual and Vioque 1996). Other studies have reported that overexpression of arginine tRNA_CCG_ from both *E. coli* (Kim et al. 1998) and *Brevibacterium albidum* (Kim et al. 1997) can complement the temperature-sensitivity caused by the *rnpA49* mutation; however, the mechanism of this has not been elucidated. Furthermore, site-directed mutagenesis of the *rnpA49* allele has identified a number of amino acid changes that can compensate the ts defect (Jovanovic et al. 2002), and additional hydroxylamine mutagenesis experiments have led to the identification of compensatory mutations in *rnpB* that can rescue the growth of A49 strains at the non-permissive temperature (Morse and Schmidt 1993). Lastly, recent work has shown that deletion of poly(A) polymerase I (PAP I, encoded by *pcnB*) can also partially rescue the ts defects caused by the *rnpA49* allele (Mohanty et al. 2020).

While these studies provide invaluable insight into RNase P assembly, structure, and function *in vivo* and demonstrate that there are many ways to rescue the *rnpA49* ts phenotype, they relied on the use of artificial approaches to examine complementation, such as plasmid overexpression, gene deletions, and mutagenesis. Subsequently, there remains a gap in the literature regarding how *E. coli* can naturally compensate for and suppress the deleterious effects brought about by the *rnpA49* mutation at non-permissive temperatures. In this work, we isolated and characterized naturally-occurring second-site suppressor mutations of the ts phenotype in an *E. coli* strain harboring the *rnpA49* allele. In agreement with past plasmid overexpression studies, we found that *E. coli* can compensate for a defective RNase P by increasing the copy number of either *rnpA49* or *rnpB* via large genome amplifications. We also identified loss of function mutations to RNase R, as well as numerous mutations impacting Lon protease expression and activity. This latter finding presents a novel link between the mutant RNase P and Lon protease, setting the stage for future investigations into RNase P regulation, degradation, and turnover.

## RESULTS AND DISCUSSION

### Isolation of suppressor mutants that enable growth of *E. coli* A49 at the non-permissive temperature

Suppressor mutants of an *E. coli* strain carrying the temperature-sensitive *rnpA49* allele were isolated from standard fluctuation assays performed at the non-permissive temperature. Briefly, approximately 10^8^ cells from 40 independent *E. coli* A49 cultures grown at permissive temperature were plated on LB-agar and incubated at the non-permissive temperature (42°C) for 48h. From these experiments, the mutation rate for *E. coli* A49 ts revertants generated by spontaneous mutation was determined to be approximately 10^-10^ to 10^-9^ per cell per generation using the bz-rates web tool (Gillet-Markowska et al. 2015).

After incubation, a single colony from each plate with growth was randomly selected, re-streaked and re-tested for growth at 42°C, and then locally sequenced to confirm the retention of the *rnpA49* allele. Seventeen isolates carrying no additional mutations to the *rnpA49* locus were selected for additional Sanger and/or whole genome sequencing to identify potential second-site suppressor mutations and for further characterization (Figure 1A; Table 1; Table S1). These suppressor isolates were subsequently classified as one of three mutation types, which are discussed in detail below: (i) gene amplifications of *rnpA49* or *rnpB*, (ii) mutations in *rnr*, and (iii) mutations in *lon*. To assess the extent of complementation, the relative growth rate of each suppressor strain was determined from growth assays in liquid medium at the permissive temperature (30°C) and at a sub-lethal temperature (40°C), to allow for comparison with the original A49 parental strain (Figure 1B; Table 1).

**Figure 1.**
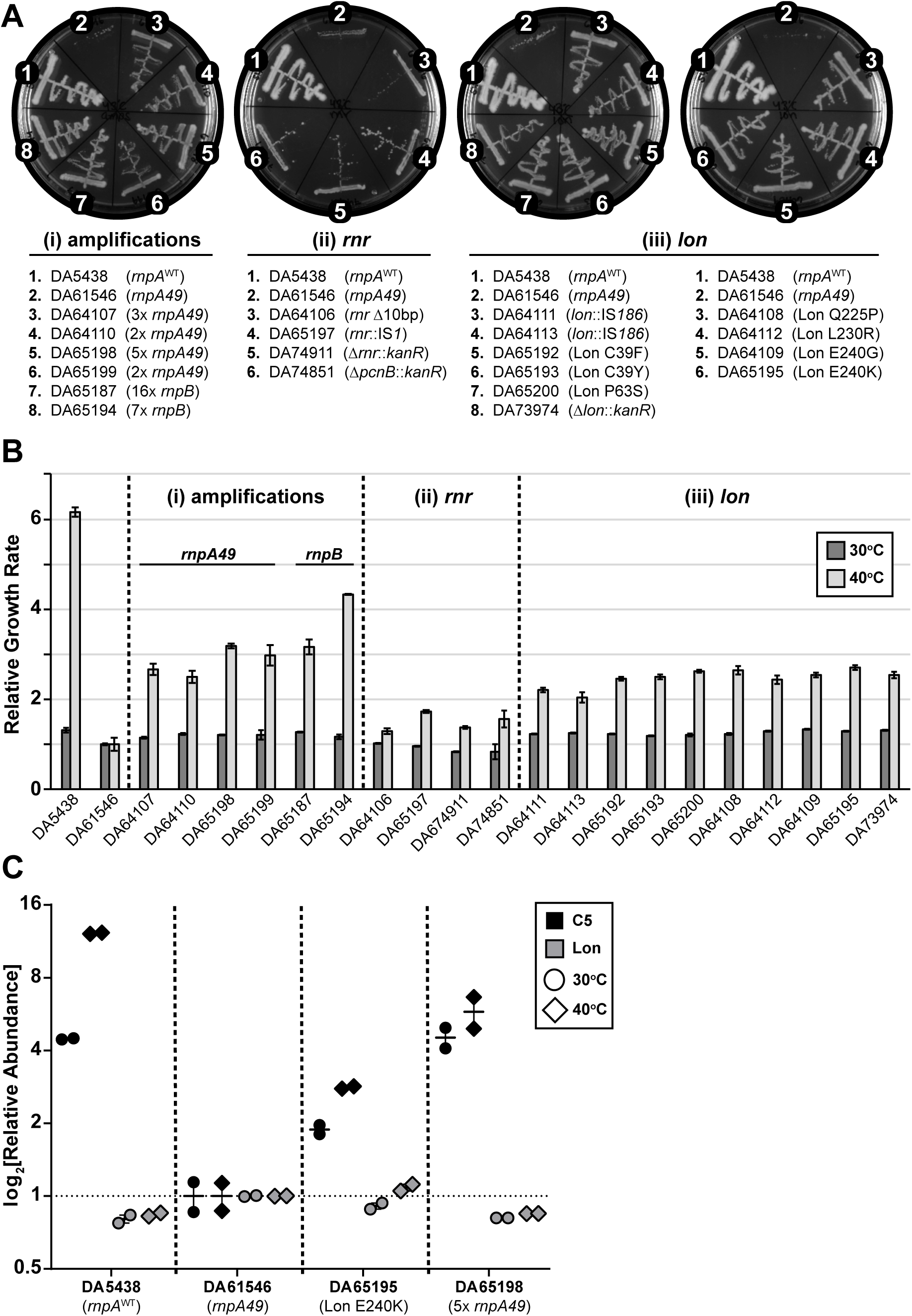
Suppressor mutations rescue the growth of *E. coli* A49 at the non-permissive temperature. (A) Growth of the *E. coli* A49 suppressor strains carrying (i) amplifications of *rnpA49* or *rnpB* or loss of function mutations to (ii) *rnr* or (iii) *lon* on LB-agar after 24h incubation at the non-permissive temperature (42°C). DA5438 is *E. coli* MG1655 carrying the wild-type *rnpA* allele; DA61546 is the *E. coli* A49 parental strain carrying the ts *rnpA49* allele. Additional control strains containing *kanR* deletion cassettes from the KEIO collection in the parental DA61546 A49 background are included for comparison. (B) Relative exponential phase growth rates of the *E. coli* A49 suppressor strains and relevant control strains as compared to the DA61546 A49 parental strain at the corresponding temperature. Growth measurements were performed in LB medium at both 30°C (permissive temperature) and 40°C (sub-lethal temperature). The values reported present the mean of at least three independent biological replicates (with two technical replicates each); error bars represent the standard error. (C) Relative abundance of either the wild-type or mutant C5 protein and Lon protease during early-to-mid exponential phase growth at the permissive (30°C) or sub-lethal temperature (40°C), as compared to the original A49 parental strain (DA61546). Strains DA61546, DA65195, and DA65198 carry the *rnpA49* mutant allele (encoding C5^A49^). Total proteome analysis was performed using two biological replicates for each strain at each temperature. Horizontal bars represent the mean between the two biological replicates; vertical I-bars represent the range. Some data points overlap.

**Table 1.**
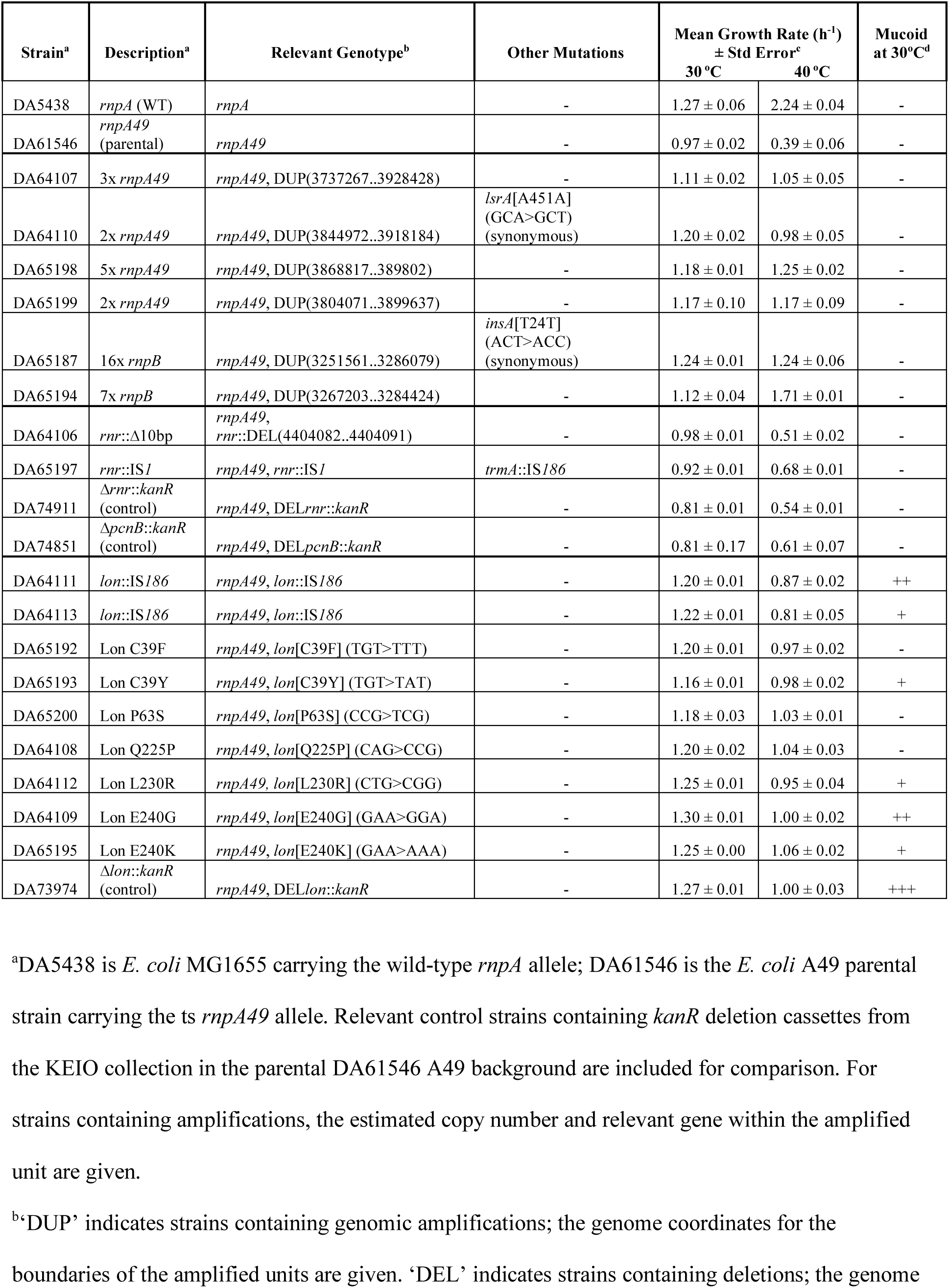
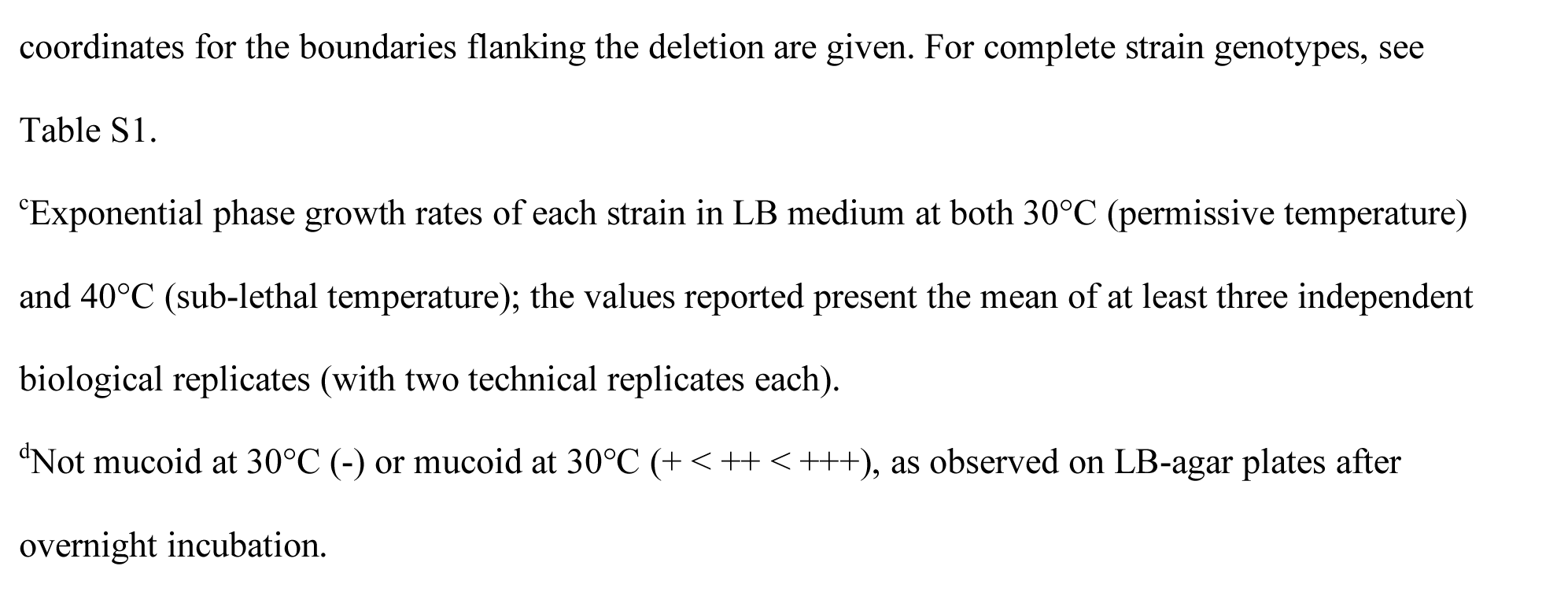
Summary of *E. coli* A49 suppressor strains isolated from this study.

| Strain<sup>a</sup> | Description<sup>a</sup> | Relevant Genotype<sup>b</sup> | Other Mutations | Mean Growth Rate (h<sup>-1</sup>)<br>± Std Error<sup>c</sup> |  | Mucoid<br>at 30°C<sup>d</sup> |
| --- | --- | --- | --- | --- | --- | --- |
|  |  |  |  | 30 °C | 40 °C |  |
| DA5438 | *rnpA* (WT) | *rnpA* | - | 1.27 ± 0.06 | 2.24 ± 0.04 | - |
| DA61546 | *rnpA49*<br>(parental) | *rnpA49* | - | 0.97 ± 0.02 | 0.39 ± 0.06 | - |
| DA64107 | 3x *rnpA49* | *rnpA49*, DUP(3737267..3928428) | - | 1.11 ± 0.02 | 1.05 ± 0.05 | - |
| DA64110 | 2x *rnpA49* | *rnpA49*, DUP(3844972..3918184) | *lsrA*[A451A]<br>(GCA>GCT)<br>(synonymous) | 1.20 ± 0.02 | 0.98 ± 0.05 | - |
| DA65198 | 5x *rnpA49* | *rnpA49*, DUP(3868817..389802) | - | 1.18 ± 0.01 | 1.25 ± 0.02 | - |
| DA65199 | 2x *rnpA49* | *rnpA49*, DUP(3804071..3899637) | - | 1.17 ± 0.10 | 1.17 ± 0.09 | - |
| DA65187 | 16x *rnpB* | *rnpA49*, DUP(3251561..3286079) | *insA*[T24T]<br>(ACT>ACC)<br>(synonymous) | 1.24 ± 0.01 | 1.24 ± 0.06 | - |
| DA65194 | 7x *rnpB* | *rnpA49*, DUP(3267203..3284424) | - | 1.12 ± 0.04 | 1.71 ± 0.01 | - |
| DA64106 | *rnr*::Δ10bp | *rnpA49*,<br>*rnr*::DEL(4404082..4404091) | - | 0.98 ± 0.01 | 0.51 ± 0.02 | - |
| DA65197 | *rnr*::IS1 | *rnpA49*, *rnr*::IS1 | *trmA*::IS186 | 0.92 ± 0.01 | 0.68 ± 0.01 | - |
| DA74911 | Δ*rnr*::*kanR*<br>(control) | *rnpA49*, DEL*rnr*::*kanR* | - | 0.81 ± 0.01 | 0.54 ± 0.01 | - |
| DA74851 | Δ*pcnB*::*kanR*<br>(control) | *rnpA49*, DEL*pcnB*::*kanR* | - | 0.81 ± 0.17 | 0.61 ± 0.07 | - |
| DA64111 | *lon*::IS186 | *rnpA49*, *lon*::IS186 | - | 1.20 ± 0.01 | 0.87 ± 0.02 | ++ |
| DA64113 | *lon*::IS186 | *rnpA49*, *lon*::IS186 | - | 1.22 ± 0.01 | 0.81 ± 0.05 | + |
| DA65192 | Lon C39F | *rnpA49*, *lon*[C39F] (TGT>TTT) | - | 1.20 ± 0.01 | 0.97 ± 0.02 | - |
| DA65193 | Lon C39Y | *rnpA49*, *lon*[C39Y] (TGT>TAT) | - | 1.16 ± 0.01 | 0.98 ± 0.02 | + |
| DA65200 | Lon P63S | *rnpA49*, *lon*[P63S] (CCG>TCG) | - | 1.18 ± 0.03 | 1.03 ± 0.01 | - |
| DA64108 | Lon Q225P | *rnpA49*, *lon*[Q225P] (CAG>CCG) | - | 1.20 ± 0.02 | 1.04 ± 0.03 | - |
| DA64112 | Lon L230R | *rnpA49*, *lon*[L230R] (CTG>CGG) | - | 1.25 ± 0.01 | 0.95 ± 0.04 | + |
| DA64109 | Lon E240G | *rnpA49*, *lon*[E240G] (GAA>GGA) | - | 1.30 ± 0.01 | 1.00 ± 0.02 | ++ |
| DA65195 | Lon E240K | *rnpA49*, *lon*[E240K] (GAA>AAA) | - | 1.25 ± 0.00 | 1.06 ± 0.02 | + |
| DA73974 | Δ*lon*::*kanR*<br>(control) | *rnpA49*, DEL*lon*::*kanR* | - | 1.27 ± 0.01 | 1.00 ± 0.03 | +++ |

<sup>a</sup>DA5438 is *E. coli* MG1655 carrying the wild-type *rnpA* allele; DA61546 is the *E. coli* A49 parental strain carrying the ts *rnpA49* allele. Relevant control strains containing *kanR* deletion cassettes from the KEIO collection in the parental DA61546 A49 background are included for comparison. For strains containing amplifications, the estimated copy number and relevant gene within the amplified unit are given.
<sup>b</sup>‘DUP’ indicates strains containing genomic amplifications; the genome coordinates for the boundaries of the amplified units are given. ‘DEL’ indicates strains containing deletions; the genome
coordinates for the boundaries flanking the deletion are given. For complete strain genotypes, see Table S1.
<sup>c</sup>Exponential phase growth rates of each strain in LB medium at both 30°C (permissive temperature) and 40°C (sub-lethal temperature); the values reported present the mean of at least three independent biological replicates (with two technical replicates each).
<sup>d</sup>Not mucoid at 30°C (-) or mucoid at 30°C (+ < ++ < +++), as observed on LB-agar plates after overnight incubation.

### Complementation via gene amplifications of *rnpA49* or *rnpB*

Of the 17 suppressor strains selected for further sequencing and study, four strains (DA64107, DA64110, DA65198, DA65199) were found to have genomic amplifications of the region containing the *rnpA49* allele, encoding the mutant C5^A49^ RNase P protein subunit, and two strains (DA65187, DA65194) were found to contain amplifications of the *rnpB* gene, encoding the M1 RNA subunit of RNase P (Table 1). The copy number of the amplified units varied among the different isolates, ranging from approximately 2-5x for *rnpA49* and 7-16x for *rnpB*, and the size of the amplified regions ranged from 29 to 192.5 kilobase pairs (kbp) for *rnpA49* and 17.2 to 34.5 kbp for *rnpB* (Table S2). Apart from a synonymous mutation in *lsrA* (predicted to encode the ATP-binding component of an AI-2 ABC transporter) that was found in one suppressor strain with a *rnpA49* amplification (DA64110) and a non-synonymous mutation in one instance of *insA* (one of the IS*1* elements on the chromosome) in a suppressor strain with a *rnpB* amplification (DA65187), no additional mutations were identified (Table 1; Table S1). As is commonly observed for genomic amplifications in bacteria, the boundaries of the majority of the amplified units were flanked by regions homologous to transposases or insertion sequences, specifically IS*1*, and in one instance, IS*4* (DA64107). However, no repeat sequences or regions of homology were identified within the break-points of the amplified units in one *rnpA49* amplification strain (DA64110), suggesting an alternate recombination mechanism not involving homologous segments (Tlsty et al. 1984; Reams and Roth 2015; Nicoloff et al. 2019) (Table S2).

In the growth assays in liquid medium, the suppressor strains containing amplifications of either *rnpA49* or *rnpB* demonstrated a slight 1.2-fold increase in growth rate relative to the original A49 parental strain at the permissive temperature. At the sub-lethal temperature, the strains carrying *rnpA49* amplifications exhibited an approximately 2.5-3-fold increase in relative growth rate, whereas the two strains with *rnpB* amplifications demonstrated a 3-4-fold increase in relative growth rate at 40°C (versus a 6-fold increase in relative growth rate for an *E. coli* strain with the wild-type *rnpA* allele at 40°C, DA5438), indicating partial complementation of the ts phenotype (Figure 1B). For each type of amplification, the strain carrying the highest copy number or level of amplification exhibited the greatest increase in relative growth rate at the sub-lethal temperature.

These findings are in agreement with past plasmid overexpression studies and confirm that increasing expression and/or gene copy number of either *rnpA49* or *rnpB* can rescue the ts defect of *E. coli* A49. As the *rnpA49* mutation is hypothesized to reduce the efficiency at which C5^A49^ assembles with the M1 RNA, overexpression of either subunit of the mutant RNase P heterodimer likely shifts the equilibrium of assembly towards formation of the RNase P holoenzyme.

The correlation between gene copy number and protein abundance was confirmed via proteomics analysis of strain DA65198. This strain carries an estimated 5x amplification of the genomic region containing the *rnpA49* allele and correspondingly exhibited a 4-5-fold increase in the abundance of the mutant C5^A49^ protein relative to the original A49 parental strain (DA61546) (Figure 1C). Interestingly, the strains carrying *rnpB* amplifications demonstrated the greatest increase in gene copy number (maximum 16x for *rnpB* versus 5x for *rnpA49*) and subsequently the highest increase in growth rate and extent of complementation. As previous work has shown that high-level overexpression of C5 is toxic in *E. coli* (Jovanovic et al. 2002), perhaps there is an upper-limit to which the *rnpA49* allele can be amplified before the toxicity of C5^A49^ overexpression outweighs complementation of the ts phenotype at the non-permissive or sub-lethal temperature. Furthermore, the sizes of the amplified units in the two *rnpB* amplification strains were on average smaller than those found in the strains containing the *rnpA49* amplifications (Table S2). It is well-established that genome amplifications inherently carry a substantial fitness cost (Nicoloff et al. 2019; Pereira et al. 2021). Thus, the smaller size of the amplified units may also contribute to the higher degree of complementation observed with the *rnpB* amplification strains. It would be interesting to see the extent to which subsequent evolution experiments would improve strain growth at the non-permissive temperature via further amplification, and/or whether the amplifications would eventually be reduced or lost upon accumulation of additional suppressor mutations.

### Complementation via loss of function mutations in *rnr*

Two suppressor strains were found to have loss of function mutations in *rnr*, encoding RNase R, a 3’ to 5’ exoribonuclease that is involved in rRNA and tmRNA maturation and turnover as well as the degradation of poly-adenylated RNAs (Andrade et al. 2009b; Condon et al. 2021). One suppressor strain (DA64106) contained a 10-base pair (bp) deletion in the N-terminus of the protein-coding sequence, resulting in a +1 frameshift after codon 59. The other strain (DA65197) contained an IS*1* insertion in *rnr*, as well as an IS*186* insertion in *trmA*, which encodes a tRNA methyltransferase that also acts a tRNA chaperone to facilitate tRNA folding (Table 1; Table S3; Figure S1).

Of all the suppressor mutants isolated, the two strains carrying *rnr* mutations exhibited the weakest complementation of the ts phenotype, with smaller colonies on plates incubated at the non-permissive temperature and slower growth rates in liquid culture (Figure 1). Both *rnr* suppressor strains grew comparable to the A49 parental strain at 30°C and exhibited only a modest 1.5-fold increase in growth rate relative to the parental A49 strain at 40°C, the sub-lethal temperature assayed.

To further confirm that loss of function mutations to *rnr* can complement the ts phenotype, we generated and assayed the growth of an A49 strain containing the /¢i*rnr*::*kanR* deletion cassette from the KEIO collection (DA74911) (Baba et al. 2006). While this deletion strain grew slower than the original A49 parental strain at the permissive temperature, it grew similarly to the two *rnr* suppressor strains recovered from our screen at the sub-lethal temperature (Figure 1B). The growth defect of the /¢i*rnr*::*kanR* control strain at 30°C is most probably due to fitness costs associated with a full deletion of *rnr* and the subsequent disruption of the genetic context surrounding the *rnr* open reading frame (ORF) and/or constitutive expression of the *kanR* cassette. Additionally, previous studies have shown that *rnr* mutants exhibit growth defects at both standard (37°C) and lower temperatures (Cairrão et al. 2003).

It is likely that the loss of function mutations to *rnr* act via the same complementation mechanism/pathway as the *pcnB* (poly(A) polymerase I, PAP I) deletion described by Mohanty et al. (2020). Because RNase P function is compromised with the *rnpA49* allele, 5’-unprocessed pre-tRNA transcripts accumulate and become substrates for polyadenylation by PAP I, which destabilizes the available pre-tRNA pool and exacerbates the ts growth defect. Consequently, deleting *pcnB* increases pre-tRNA levels, ultimately improving the processing and/or aminoacylation of the premature tRNAs to facilitate survival at the non-permissive temperature (Mohanty et al. 2020). As RNase R targets poly-adenylated RNAs for degradation, and *rnr* expression is elevated at higher temperatures (Andrade et al. 2009a), it likely further destabilizes the PAP I poly-adenylated pre-tRNAs at 42°C. Therefore, inactivating and/or deleting *rnr* also increases the pre-tRNA pool available for aminoacylation and subsequent survival at the non-permissive temperature.

This is further supported by the fact that our A49 parental strain containing the ϕ..*pcnB*::*kanR* deletion cassette from the KEIO collection (DA74851) also behaves similar to both the naturally-occurring *rnr* suppressor mutants isolated from our screen and the constructed A49 ϕ..*rnr*::*kanR* control strain at the sub-lethal temperature (Figure 1). Like the ϕ..*rnr*::*kanR* strain, the ϕ..*pcnB*::*kanR* strain also exhibits a slight growth defect at the permissive temperature relative to the parental A49 strain. Again, this is likely due to costs associated with *kanR* expression and/or potential disruption of the regulatory sequences surrounding or overlapping the fully-deleted *pcnB* ORF.

Alternatively, it is also possible that RNase R is involved in the turnover and degradation of M1 RNA. Thus, inactivation of *rnr* may directly improve the assembly and stability of the mutant RNase P holoenzyme at the non-permissive temperature. However, follow-up experiments are necessary to confirm this. While RNase E is involved in M1 RNA maturation (Lundberg and Altman 1995; Ko et al. 2008), and the M1 RNA is likely stable for extended periods of time (Jain et al. 1982), the enzymes and pathways responsible for M1 RNA turnover and degradation are still unknown.

### Complementation via promoter disruption and nonsynonymous mutations in *lon*

The most common type of mutations recovered from our fluctuation assays targeted *lon*, encoding the ATP-dependent AAA+ protease, Lon. In bacteria, Lon is a global regulator that coordinates a number of important processes such as stress response, DNA replication and repair, and virulence via the degradation of misfolded proteins, rapid turnover of key regulatory proteins, and in some instances, acting as a protein-folding chaperone (Tsilibaris et al. 2006; Van Melderen and Aertsen 2009; Gur 2013). In *E. coli*, Lon subunits equilibrate between hexamers and dodecamers under physiological conditions (Wohlever et al. 2013). Two strains were found to have an IS*186* insertion in the *lon* promoter region and seven strains were found to contain SNPs resulting in nonsynonymous mutations in the *lon* protein-coding region. Six of the nine *lon* mutant suppressor strains were mucoid when grown at 30°C (DA64109, DA64111, DA64112, DA64113, DA65193, DA65195), a characteristic phenotype of *E. coli lon* mutants (Bush and Markovitz 1973) (Table 1).

Previous studies have shown that the *lon* promoter is a hot spot for IS*186* insertion and that IS*186*-mediated *lon* promoter disruption is a fairly frequent mutation that inhibits *lon* expression under specific selective conditions, such as in the presence of certain antibiotics (SaiSree et al. 2001; Nicoloff et al. 2007; Nicoloff and Andersson 2013) (Table S3). The *lon*::IS*186* suppressor strains demonstrated an approximately 1.3-fold increase in relative growth rate at 30°C, and a 2-fold increase in relative growth rate at the sub-lethal 40°C when compared to the original A49 parental strain (Figure 1B).

Similarly, the suppressor strains carrying the nonsynonymous changes to the *lon* protein-coding sequence also exhibited a 1.3-fold increase in relative growth rate at the permissive temperature. However, these strains complemented the ts defect slightly better than the *lon*::IS*186* suppressor strains, with an average 2.6-fold increase in relative growth rate at the sub-lethal temperature. Of the seven *lon* protein-coding mutants isolated, four strains contained substitutions at or near residue E240 (Table 1). The E240K mutation has previously been shown to favor Lon’s dodecamer conformation and affects substrate recognition and degradation activity. Furthermore, the region surrounding residue 240 has been implicated in substrate recognition and activation of Lon’s protease and ATPase activity (Ebel et al. 1999; Cheng et al. 2012; Wohlever et al. 2013). This suggests that the suppressor mutants carrying the single nonsynonymous mutations within this region (DA64108: Q225P, DA64112: L230R, DA64109: E240G, DA65195: E240K) likely have a partial or complete loss of Lon function. It is less clear how the suppressor mutations located closer to the N-terminus affect Lon function and/or expression (DA65192: C39F, DA65193: C39Y, DA65200: P63S). Nevertheless, these three suppressor strains behave similar to the *lon* suppressor strains that carry mutations known to alter Lon expression and/or function, and the suppressor strain with the C39Y mutation (DA65193) is mucoid at 30°C, indicating that nonsynonymous mutations close to the N-terminus can indeed impact Lon activity and/or expression. Lastly, a control A49 strain containing the /-.*lon*::*kanR* deletion cassette from the KEIO collection (DA73974) was constructed and assayed, confirming that loss of Lon function can partially complement the ts defect caused by the *rnpA49* allele (Figure 1).

To our knowledge, the wild-type C5 protein subunit of RNase P has not yet been identified as a substrate for Lon protease (Heuveling et al. 2008), and the observation that wild-type C5 protein abundance is similar in *lon*+ (DA5438) and *lon*-(DA22599) *E. coli* MG1655 strains supports this (Figure S2). However, it is possible that the mutant C5^A49^ is degraded by Lon. Thus, *lon* loss of function mutations would stabilize C5^A49^ and the overall mutant RNase P holoenzyme to partially suppress the *rnpA49* temperature-sensitivity. This is consistent with findings from a similar study in which a mutant RNA polymerase sigma subunit was found to be a direct substrate for Lon proteolysis and *lon* loss of function mutations suppressed the ts phenotype caused by the mutant *rpoD800* allele (Grossman et al. 1983).

To investigate how loss of function mutations to Lon protease impact the relative abundance of both the wild-type and mutant C5 protein, we performed total proteome analysis on select *E. coli* strains during early-to-mid exponential phase growth at both the permissive (30°C) and sub-lethal temperatures (40°C) (Figure 1C; Figure S2). While the relative abundance of the mutant Lon[E240K] in suppressor strain DA65195 remained similar to that of both the A49 parental strain (DA61546) and the MG1655 wild-type strain (DA5438), suppressor strain DA65195 demonstrated an approximately 2-3-fold increase in the relative abundance of the mutant C5^A49^ protein subunit as compared to the original ts A49 parental strain (DA61546). This increase in C5^A49^ abundance coincides with the almost 3-fold increase in the suppressor strain’s relative growth rate at the non-permissive temperature and indicates that the mutant C5^A49^ protein is potentially a substrate of Lon protease (Figure 1B). It is also possible that Lon indirectly impacts C5^A49^ stability via the regulation of other proteases and/or pathways. Nevertheless, the enzymes and pathways involved in the degradation and turnover of RNase P remain unknown, and additional studies beyond the scope of this work are required to further elucidate the pathways and mechanisms involved in regulating RNase P levels.

## CONCLUDING REMARKS

In this study, we report for the first time naturally-occurring second-site compensatory mutations of the *rnpA49* ts phenotype in *E. coli*. Although a relatively small screen, similar mutations were recovered multiple independent times across the 17 suppressor strains isolated, suggesting that these are frequent targets for compensatory evolution. The compensatory mechanisms underlying the genomic amplifications of the regions containing the genes encoding either subunit of the mutant RNase P complex are well-established from previous plasmid overexpression studies, and there is precedent for the role of RNase R loss of function mutations in the literature. However, follow-up studies are necessary to determine the exact mechanism behind the *rnr* suppressor mutations. Additionally, we found that *E. coli* frequently suppresses the ts defect caused by the *rnpA49* allele via mutations that likely inactivate or alter the activity or expression of Lon protease and that select suppressor strains with *lon* mutations have increased C5^A49^ abundance. This suggests that the mutant C5^A49^ protein is potentially a substrate for Lon degradation and/or Lon protease is indirectly involved in the regulation of C5^A49^ levels.

It is interesting to note that while the isolated strains grow comparable to wild-type *E. coli* (DA5438) at the permissive temperature, the suppressor mutations only partially complement the ts phenotype at the sub-lethal and non-permissive temperatures, and growth rates are not restored to wild-type levels at the higher temperatures. This can potentially be attributed to the inherent costs of the suppressor mutations themselves. As mentioned previously, large genome amplifications can have substantial impacts on cell fitness (Nicoloff et al. 2019; Pereira et al. 2021), high-level C5 overexpression is known to be toxic (Jovanovic et al. 2002), and the disruption of the various regulatory functions of RNase R and Lon likely have off-target effects on overall fitness. Furthermore, only single relevant suppressor mutations were identified in each strain. Therefore, it is possible that additional mutations could arise with subsequent evolution experiments to more fully complement the ts defect and/or compensate for the fitness costs brought about by the original suppressor mutations.

Apart from the overexpression of either RNase P subunit via genome amplifications, the other second-site suppressor mutations recovered were loss of function mutations in other enzymes, suggesting that there is no promiscuous and/or moonlighting activity from alternative *E. coli* RNases/proteins that can compensate for the mutant RNase P. To determine if the overexpression of other *E. coli* ORFs can in fact rescue the *rnpA49* ts defect, we introduced a plasmid library containing pooled clones from the ASKA collection (representing 4,000 cloned ORFs from *E. coli*) into our *E. coli* A49 strain and screened for ORFs that could enable strain growth at the non-permissive temperature (42°C) (Kitagawa et al. 2005). The gene encoding the wild-type C5 protein, *rnpA*, was the only “positive hit” recovered from this screen, further supporting our observation that complementation is primarily driven by overexpression of RNase P or loss of function mutations in other genes, rather than via gain of function mutations and/or increased expression of other potentially promiscuous enzymes (Figure S3).

For decades, the molecular, biochemical, and structural elements governing RNase P function have been the subject of hundreds of studies. Nevertheless, the regulation of RNase P expression, its turnover and degradation, and the mechanisms underlying certain RNase P mutations are still not fully understood. While informative, the previous efforts to examine the suppression of various RNase P mutant phenotypes were limited to “artificial” plasmid overexpression, gene deletion, and mutagenesis approaches. Our work complements these past studies and describes ways by which *E. coli* A49 naturally compensates for the RNase P assembly and fitness defects caused by the mutant *rnpA49* allele. All of the suppressor mutations isolated from our study appear to drive equilibrium to favor the formation and stability of the mutant RNase P holoenzyme (*rnpA49* and *rnpB* amplifications, *lon* mutations) or the stability of the enzyme’s pre-tRNA substrates (*rnr* mutations) to ultimately improve RNase P function and cell survival at the non-permissive temperature. Our findings suggest novel links between RNase P and the regulatory pathways involving RNase R and Lon protease, and implicate the mutant C5^A49^ RNase P protein subunit as a target for Lon proteolysis. We foresee that this work will lay the groundwork for future investigations into RNase P regulation and degradation.

## MATERIALS AND METHODS

### Bacterial strains and growth conditions

The temperature-sensitive *rnpA49* allele was transferred from the original mutagenized *E. coli* A49 strain (*E. coli* Genetic Stock Center Strain #: 6465; Strain Designation: N2020; F-, *lacZ8*(Am), *trpA36*, *glyA34*, *argA52*, *rpsL999*(*strR*), *rnpA49*(ts), *ilvG866*(Act)) (Apirion 1980) to an *E. coli* MG1655 background (F-, lambda-, *rph-1*) using P1 transduction (Thomason et al. 2007). Presence of the temperature-sensitive mutation of interest was confirmed via lack of growth at the non-permissive temperature (42°C) and local and whole genome sequencing (MG1655 A49 strain designated as DA61546). For all experiments, strains were grown in lysogeny broth (LB Miller; 10 g/l NaCl, 10 g/l tryptone, 5 g/l yeast extract; Sigma-Aldrich) or plated on LB supplemented with 1.5% (w/v) agar (LA; Millipore). All strains generated in this study were cryo-preserved at –80°C in LB supplemented with 10% DMSO. A49 strains containing additional gene deletions were constructed by P1 transduction of the *kanR* cassette from the corresponding KEIO collection strain and selection on LA supplemented with 50 μg/ml kanamycin at 30°C (Baba et al. 2006; Thomason et al. 2007). See Table S1 for the list of strains used, isolated, and/or constructed for this study.

### Fluctuation assays

Twenty cultures of *E.coli* A49 strain DA61546 inoculated from independent colonies were grown overnight in 1 ml LB at 30°C with shaking at 190 rpm. For the fluctuation assay, approximately 10^8^ cells (50 μl) from each overnight culture were plated on LA prewarmed to 42°C, and then incubated at 42°C (non-permissive temperature) for a total of 48h. Plates were monitored and colonies were counted after 16h, 20h, and 48h incubation. After 48h, a single colony was randomly chosen from each plate with growth, re-isolated on LA plates incubated at 30°C, re-tested for growth at 42°C, and saved for future studies. To confirm the amount of cells plated, serial dilutions of each overnight culture were plated on LA, incubated at 30°C overnight, and colonies were counted. Mutation rate was determined using the bz-rates webtool, which implements the Ma-Sandri-Sakar maximum likelihood estimator (Hamon and Ycart 2012; Gillet-Markowska et al. 2015). This experiment was performed two independent times, for a total of 40 independent colonies from A49 strain DA61546 being assayed.

### Whole genome sequencing

Prior to whole genome sequencing, the *rnpA49* and *lon* loci of select ts revertant strains isolated from the fluctuation assays were PCR-amplified and Sanger sequenced (Eurofins) to confirm presence of the *rnpA49* and wild-type *lon* alleles (see Table S4 for the list of oligos used for PCR and local sequencing). These strains were then grown for whole genome sequencing in 1 ml LB overnight at 30°C with shaking at 190 rpm. Genomic DNA was extracted from 0.5 ml of each culture using an Epicentre MasterPure Complete DNA and RNA Purification Kit according to the manufacturer’s protocol. DNA was resuspended in water, concentrations were determined using a Nanodrop 1000 (Thermo Scientific) and Qubit 2.0 fluorometer (Invitrogen, DNA Broad-Range kit), and sequencing was performed in-house using an Illumina MiSeq platform with a Nextera XT DNA library preparation kit. Sequences were mapped to the reference genomes and mutations were identified using CLC Genomic Workbench software (Qiagen). The extent of DNA amplification was estimated by dividing the average sequence coverage of the amplifications by the average sequence coverage of the rest of the chromosome in CLC Genomic Workbench (Qiagen) as described previously (Nicoloff et al. 2019). All raw sequence reads for the whole genome sequencing from this study are deposited in the Sequence Read Archive (SRA) at the National Center for Biotechnology Information (NCBI) under BioProject Accession PRJNA1032291.

### ASKA library screen

A plasmid library (representing 4,000 cloned ORFs from *E. coli*) from approximately 460,000 pooled clones from the ASKA collection (Kitagawa et al. 2005) was isolated using an EZNA Plasmid DNA Mini Kit I (Omega Bio-Tek). Electrocompetent *E. coli* A49 strain DA61546 cells were prepared as described previously, but with incubation at 30°C (Knopp et al. 2019), and approximately 300 ng of the ASKA plasmid library or the empty pCA24N ASKA plasmid were transformed into 40 μl cells via electroporation. Cells were immediately recovered in 1 ml LB at 30°C with shaking at 190 rpm for 1h. After incubation, a dilution series for each transformation was plated on LA supplemented with 20 μg/ml chloramphenicol, which was then incubated at 30°C overnight, and the colonies were counted to determine the transformation efficiency and subsequently estimate the diversity of the library covered (approx. 10^6^ total transformants). The remaining transformant recovery cultures were plated on LA supplemented with 20 μg/ml chloramphenicol and 1 mM IPTG and allowed to incubate at 42°C for approx. 30h. Any colonies that appeared were re-streaked on LA supplemented with 20 μg/ml chloramphenicol and 1 mM IPTG and incubated at 42°C overnight to confirm rescue of the ts phenotype. Plasmids were isolated from these “positive hits” using the EZNA Plasmid DNA Mini Kit I (Omega Bio-Tek), sequenced using Sanger sequencing (Eurofins) to identify the cloned ORF recovered (Table S4), re-transformed into DA61546, and re-tested for growth at the non-permissive temperature as described above.

### Growth assays

Growth curves were obtained using a Bioscreen C instrument (Oy Growth Curves AB, Ltd., Finland). Overnight cultures (1 ml LB) of select strains were grown in triplicate from independent colonies at 30°C with shaking at 190 rpm and used to inoculate 1 ml fresh LB cultures (1 μl, 1:1000 dilution). From this dilution, two 300 μl aliquots were transferred to Bioscreen C honeycomb plates to serve as technical replicates. Plates were incubated in the Bioscreen C for 24h (30°C assays, permissive temperature) or 4 days (40°C assays, sub-lethal temperature) with shaking and the OD_600_ was measured every 4 min. The background OD_600_ measurements from wells containing blank, un-inoculated medium were subtracted from the OD600 measurements. Growth rates were calculated from the linear slope of ln(OD_600_) during exponential phase (ln(OD_600_) -3.5 to -2.5), and relative growth rate was calculated by the dividing the growth rate of each suppressor or control strain by that of the parental A49 strain (DA61546) in the corresponding condition. The reported values represent the mean of at least three biological replicates, with two technical replicates each; the reported error is the standard error.

### Proteomics sample preparation

Cultures (5 ml LB) started from two independent colonies from select suppressor and control strains were grown overnight at 30°C with shaking (190 rpm). These were used to inoculate two 500 ml flasks containing 60 ml LB to a starting OD_600_ of 0.06. For each strain/biological replicate, one flask was incubated at 30°C with shaking (190 rpm) and one flask was incubated at 40°C (water bath) with shaking (175 rpm), until an OD_600_ ∼0.2-0.25 was reached. Flasks were then cooled on ice for 10-20 mins, cells were pelleted at 4500 rpm, 4°C for 15 mins, washed twice with 1 ml phosphate-buffered saline (PBS; 8 g/l NaCl, 0.2 g/l KCl, 1.44 g/l Na_2_HPO_4_, and 0.24 g/l KH_2_PO_4_), and then stored at – 80°C until proteomics experiments were performed.

The bacterial samples were homogenized using a FastPrep-24 instrument (MP Biomedicals) with Lysing Matrix B in 4% SDS, 1 mM DTT in 50 mM HEPES [pH 7], and protein concentration was determined using the Pierce BCA Protein Assay Kit (Thermo Scientific) on a SpectraMax iD3 (Molecular Devices). Aliquots of 200 µg protein were reduced in 10 mM dithiothreitol at 56°C for 30 min, then alkylated with 20 mM chloroacetamide at room temperature for 10 min. Protein samples were added to washed hydrophobic and hydrophilic Sera-Mag™ SpeedBeads (Carboxylate-Modified, Cytiva) in a bead-to-protein ratio of 10:1. The SP3-workflow was adapted from the protein and peptide clean-up for mass spectrometry protocol provided by the manufacturer. In short, proteins were precipitated on the beads by acetonitrile, washed with ethanol, and dried at room temperature. For digestion, 50 mM TEAB and 2 µg LysC+trypsin (Promega) were added, and incubation took place overnight at 37°C while shaking. An additional portion of 2 µg trypsin (Thermo Fisher Scientific) was added and digested for 3h. The peptide concentration was determined using the Pierce™ Quantitative Peptide Assay (Thermo Scientific). Aliquots of 30 µg peptides were labelled using TMTpro 18-plex isobaric mass tagging reagents (Thermo Fisher Scientific) according to the manufacturer’s instructions. All samples were pooled into one TMT-set, and peptides were purified using HiPPR detergent removal kit and Pierce peptide desalting spin columns (both Thermo Fisher Scientific), according to the manufacturer’s instructions. The TMT-sets were fractionated by basic reversed-phase chromatography using a Dionex Ultimate 3000 UPLC system (Thermo Fisher Scientific). Peptide separations were performed using a reversed-phase XBridge BEH C18 column (3.5 µm, 3.0x250 mm, Waters Corporation) and a stepped gradient from 3% to 54% solvent B over 65 min followed by an increase to 80% solvent B at a flow of 200 µL/min. Solvent A was 25% ammonia and solvent B was 84% acetonitrile. The 96 primary fractions were combined into 15 final fractions which were evaporated and reconstituted in 3% acetonitrile, 0.1% trifluoroacetic acid for LC-MS3 analysis.

### LC-MS3 analysis

The fractions were analyzed on an Orbitrap Eclipse Tribrid mass spectrometer equipped with the FAIMS Pro ion mobility system and interfaced with an Easy-nLC1200 liquid chromatography system (both Thermo Fisher Scientific). Peptides were trapped on an Acclaim Pepmap 100 C18 trap column (100 µm x 2 cm, particle size 5 µm, Thermo Fisher Scientific) and separated on an in-house packed analytical column (35 cm x 75 µm, particle size 3 µm, Reprosil-Pur C18, Dr. Maisch) using a stepped gradient from 4% to 80% acetonitrile in 0.2% formic acid over 75 min at a flow of 300 nL/min. FAIMS Pro was alternating between the compensation voltages (CV) of -50 and -70, and the same data-dependent settings were used at both CVs. The precursor ion mass spectra were acquired at a resolution of 120,000 and an m/z range of 375-1500. Using a cycle time of 1.5 seconds, the most abundant precursors with charges 2–7 were isolated with an m/z window of 0.7 and fragmented by collision induced dissociation (CID) at 30%. Fragment spectra were recorded in the ion trap at Rapid scan rate. The ten most abundant MS2 fragment ions were isolated using multi-notch isolation for further MS3 fractionation. MS3 fractionation was performed using higher-energy collision dissociation (HCD) at 55% and the MS3 spectra were recorded in the Orbitrap at 50000 resolution and an m/z range of 100–500.

### Proteomic data analysis

Identification and relative quantification were performed using Proteome Discoverer version 2.4 (Thermo Fisher Scientific). The data was matched against *E.coli* SwissProt database (5385 entries, May 2023). Database matching was performed using Sequest as a search engine with a precursor tolerance of 5 ppm and a fragment ion tolerance of 0.6 Da. Tryptic peptides were accepted with 1 missed cleavage; methionine oxidation was set as a variable modification and cysteine carbamidomethylation, TMTpro on lysine and peptide N-termini were set as fixed modifications. Percolator was used for PSM validation with a strict FDR threshold of 1%. For quantification, TMT reporter ions were identified in the MS3 HCD spectra with 3 mmu mass tolerance and the TMT reporter intensity values for each sample were normalized on the total peptide amount. SPS Mass Match threshold was set to 65%; a Sequest XCorr threshold score of 2 was chosen. Only unique peptides were used for relative quantification and proteins were required to pass a protein FDR of 5%. The mass spectrometry proteomics data have been deposited to the ProteomeXchange Consortium via the PRIDE partner repository with the dataset identifier PXD047498 (Perez-Riverol et al. 2019).

## ACKNOWLEDGEMENTS

This work was supported by grants from the Wallenberg Foundation (grant 2018.0168 to DIA) and the Swedish Research Council (grant 2019-02091 to DIA). The funders had no role in study design, data collection and analysis, decision to publish, or preparation of the manuscript. The quantitative proteomics analysis was performed by Evelin Berger at the Proteomics Core Facility, Sahlgrenska academy, Gothenburg University, with financial support from SciLifeLab and BioMS. The authors thank Omar Mahmud Warsi and Hervé Nicoloff for their assistance with whole genome sequencing and analysis, Roderich Römhild for help with growth curve analysis, and Nikolaos Fatsis-Kavalopoulos for help with data visualization.

## SUPPLEMENTAL MATERIALS (attached separately)

**Table S1. *Escherichia coli* strains used in this study.**

**Table S2. Summary of *Escherichia coli* A49 suppressor strains containing genomic amplifications.**

**Table S3. Summary of *Escherichia coli* A49 suppressor strains containing insertions.**

**Table S4. Oligonucleotides used in this study.**

**Figure S1. Disruption of *trmA* does not contribute to the rescue phenotype of suppressor strain DA65197.**

**Figure S2. Deleting *lon* does not affect wild-type C5 protein abundance during early-to-mid exponential phase growth.**

**Figure S3. Only overexpression of *rnpA*^WT^ was found to rescue *E. coli* strain A49 following an ASKA library screen.**

## REFERENCES

1. Agrawal A, Mohanty BK, Kushner SR. 2014. Processing of the seven valine tRNAs in Escherichia coli involves novel features of RNase P. Nucleic Acids Res 42: 11166–11179.

2. Alifano P, Rivellini F, Piscitelli C, Arraiano CM, Bruni CB, Carlomagno MS. 1994. Ribonuclease E provides substrates for ribonuclease P-dependent processing of a polycistronic mRNA. Genes Dev 8: 3021–3031.

3. Altman S, Kirsebom LA. 1999. Ribonuclease P. In The RNA World (eds. R.F. Gesteland, T. Cech, and J.F. Atkins), pp. 351–380, Cold Spring Harbor Laboratory Press, Cold Spring Harbor, NY.

4. Andrade JM, Hajnsdorf E, Régnier P, Arraiano CM. 2009a. The poly(A)-dependent degradation pathway of rpsO mRNA is primarily mediated by RNase R. RNA 15: 316–326.

5. Andrade JM, Pobre V, Silva IJ, Domingues S, Arraiano CM. 2009b. The Role of 3′-5′ Exoribonucleases in RNA Degradation. Prog Mol Biol Transl Sci 85: 187–229.

6. Apirion D. 1980. Genetic mapping and some characterization of the rnpA49 mutation of Escherichia coli that affects the RNA-processing enzyme ribonuclease P. Genetics 94: 291–299.

7. Baba T, Ara T, Hasegawa M, Takai Y, Okumura Y, Baba M, Datsenko KA, Tomita M, Wanner BL, Mori H. 2006. Construction of Escherichia coli K-12 in-frame, single-gene knockout mutants: the Keio collection. Mol Syst Biol 2: 2006 0008.

8. Baer MF, Wesolowski D, Altman S. 1989. Characterization In Vitro of the Defect in a Temperature-Sensitive Mutant of the Protein Subunit of RNase P from Escherichia coli. J Bacteriol 171: 6862–6866.

9. Bothwell ALM, Garber RL, Altman S. 1976. Nucleotide sequence and in vitro processing of a precursor molecule to Escherichia coli 4.5 S RNA. J Biol Chem 251: 7709–7716.

10. Buck AH, Dalby AB, Poole AW, Kazantsev A V., Pace NR. 2005. Protein activation of a ribozyme: The role of bacterial RNase P protein. EMBO J 24: 3360–3368.

11. Bush JW, Markovitz A. 1973. The Genetic Basis for Mucoidy and Radiation Sensitivity. Genetics 74: 215–225.

12. Cairrão F, Cruz A, Mori H, Arraiano CM. 2003. Cold shock induction of RNase R and its role in the maturation of the quality control mediator SsrA/tmRNA. Mol Microbiol 50: 1349–1360.

13. Cheng I, Mikita N, Fishovitz J, Frase H, Wintrode P, Lee I. 2012. Identification of a region in the N-terminus of Escherichia coli Lon that affects ATPase, substrate translocation and proteolytic activity. J Mol Biol 418: 208–225.

14. Condon C, Pellegrini O, Gilet L, Durand S, Braun F. 2021. Walking from E. coli to B. subtilis, one ribonuclease at a time. Comptes Rendus - Biol 344: 357–371.

15. Ebel W, Skinner MM, Dierksen KP, Scott JM, Trempy JE. 1999. A conserved domain in Escherichia coli Lon protease is involved in substrate discriminator activity. J Bacteriol 181: 2236–2243.

16. Gillet-Markowska A, Louvel G, Fischer G. 2015. bz-rates: A web tool to estimate mutation rates from fluctuation analysis. G3 *Genes, Genomes, Genet* **5**: 2323–2327.

17. Grossman AD, Burgess RR, Walter W, Gross CA. 1983. Mutations in the Lon gene of E. coli K12 phenotypically suppress a mutation in the sigma subunit of RNA polymerase. Cell 32: 151–159.

17a. Guerrier-Takada C, Gardiner K, Marsh T, Pace N, Altman S. 1983. The RNA moiety of ribonuclease P is the catalytic subunit of the enzyme. Cell 35: 849–857.

18. Gur E. 2013. The Lon AAA+ protease. In Regulated Proteolysis in Microorganisms (ed. D.A. Dougan), Vol. 66 of, pp. 35–51, Springer, New York.

19. Hamon A, Ycart B. 2012. Statistics for the Luria-Delbrück distribution. Electron J Stat 6: 1251–1272.

20. Harris ME, Frank D, Pace NR. 1998. Structure and catalytic function of the bacterial ribonuclease P ribozyme. In RNA Structure and Function (eds. R.W. Simons and M. Grunberg-Manago), pp. 309–337, Cold Spring Harbor Laboratory Press, Cold Spring Harbor, NY.

21. Heuveling J, Possling A, Hengge R. 2008. A role for Lon protease in the control of the acid resistance genes of Escherichia coli. Mol Microbiol 69: 534–547.

22. Jain SK, Gurevitz M, Apirion D. 1982. A small RNA that complements mutants in the RNA processing enzyme ribonuclease P. J Mol Biol 162: 515–533.

23. Jovanovic M, Sanchez R, Altman S, Gopalan V. 2002. Elucidation of structure-function relationships in the protein subunit of bacterial RNase P using a genetic complementation approach. Nucleic Acids Res 30: 5065–5073.

24. Kim MS, Kim S, Kim SC, Lee YM, Jeon ES, Park CU, Lee Y. 1997. The Brevibacterium albidum gene encoding the arginine tRNA(CCG) complements the growth defect of an Escherichia coli strain carrying a thermosensitive mutation in the rnpA gene at the nonpermissive temperature. Mol Gen Genet 254: 464–468.

25. Kim MS, Park BH, Kim S, Lee YJ, Chung JH, Lee Y. 1998. Complementation of the growth defect of an rnpA49 mutant of Escherichia coli by overexpression of arginine tRNA(CCG). Biochem Mol Biol Int 46: 1153–1160.

26. Kirsebom LA, Baer MF, Altman S. 1988. Differential effects of mutations in the protein and RNA moieties of RNase P on the efficiency of suppression by various tRNA suppressors. J Mol Biol 204: 879–888.

27. Kirsebom LA, Trobro S. 2009. RNase P RNA-mediated cleavage. IUBMB Life 61: 189–200.

28. Kitagawa M, Ara T, Arifuzzaman M, Ioka-Nakamichi T, Inamoto E, Toyonaga H, Mori H. 2005. Complete set of ORF clones of Escherichia coli ASKA library (A complete set of E. coli K-12 ORF archive): unique resources for biological research. DNA Res 12: 291–299.

29. Knopp M, Gudmundsdottir JS, Nilsson T, Konig F, Warsi O, Rajer F, Adelroth P, Andersson DI. 2019. De Novo Emergence of Peptides That Confer Antibiotic Resistance. MBio 10: e00837–19.

30. Ko JH, Han K, Kim Y, Sim S, Kim KS, Lee SJ, Cho B, Lee K, Lee Y. 2008. Dual function of RNase E for control of M1 RNA biosynthesis in Escherichia coli. Biochemistry 47: 762–770.

31. Komine Y, Kitabatake M, Yokogawa T, Nishikawa K, Inokuchi H. 1994. A tRNA-like structure is present in 10Sa RNA, a small stable RNA from Escherichia coli. Proc Natl Acad Sci U S A 91: 9223–9227.

32. Li Y, Altman S. 2003. A specific endoribonuclease, RNase P, affects gene expression of polycistronic operon mRNAs. Proc Natl Acad Sci U S A 100: 13213–13218.

33. Li Y, Cole K, Altman S. 2003. The effect of a single, temperature-sensitive mutation on global gene expression in Escherichia coli. RNA 9: 518–532.

34. Lin HC, Zhao J, Niland CN, Tran B, Jankowsky E, Harris ME. 2016. Analysis of the RNA Binding Specificity Landscape of C5 Protein Reveals Structure and Sequence Preferences that Direct RNase P Specificity. Cell Chem Biol 23: 1271–1281.

35. Lundberg U, Altman S. 1995. Processing of the precursor to the catalytic RNA subunit of RNase P from Escherichia coli. RNA 1: 327–334.

36. Mohanty BK, Agrawal A, Kushner SR. 2020. Generation of pre-tRNAs from polycistronic operons is the essential function of RNase P in Escherichia coli. Nucleic Acids Res 48: 2564–2578.

37. Mohanty BK, Kushner SR. 2008. Rho-independent transcription terminators inhibit RNase P processing of the secG leuU and metT tRNA polycistronic transcripts in Escherichia coli. Nucleic Acids Res 36: 364–375.

38. Mohanty BK, Kushner SR. 2007. Ribonuclease P processes polycistronic tRNA transcripts in Escherichia coli independent of ribonuclease E. Nucleic Acids Res 35: 7614–7625.

39. Morse DP, Schmidt FJ. 1992. Sequences encoding the protein and RNA components of ribonuclease P from Streptomyces bikiniensis var. zorbonensis. Gene 117: 61–66.

40. Morse DP, Schmidt FJ. 1993. Suppression of loss-of-function mutations in Escherichia coli Ribonucelase P RNA (M1 RNA) by a specific base-pair disruption. J Mol Biol 230: 11–14.

41. Motamedi H, Lee Y, Schmidt FJ. 1984. Tandem promoters preceding the gene for the M1 RNA component of Escherichia coli ribonuclease P. Proc Natl Acad Sci U S A 81: 3959–3963.

42. Nicoloff H, Andersson DI. 2013. Lon protease inactivation, or translocation of the lon gene, potentiate bacterial evolution to antibiotic resistance. Mol Microbiol 90: 1233–1248.

43. Nicoloff H, Hjort K, Levin BR, Andersson DI. 2019. The high prevalence of antibiotic heteroresistance in pathogenic bacteria is mainly caused by gene amplification. Nat Microbiol 4: 504–514.

44. Nicoloff H, Perreten V, Levy SB. 2007. Increased genome instability in Escherichia coli lon mutants: Relation to emergence of multiple-antibiotic-resistant (Mar) mutants caused by insertion sequence elements and large tandem genomic amplifications. Antimicrob Agents Chemother 51: 1293–1303.

45. Pascual A, Vioque A. 1996. Cloning, purification and characterization of the protein subunit of ribonuclease P from the cyanobacterium Synechocystic sp. PCC 6803. *Eur J Biochem* **241**: 17– 24.

46. Pereira C, Larsson J, Hjort K, Elf J, Andersson DI. 2021. The highly dynamic nature of bacterial heteroresistance impairs its clinical detection. Commun Biol 4: 1–12.

47. Perez-Riverol Y, Csordas A, Bai J, Bernal-Llinares M, Hewapathirana S, Kundu DJ, Inuganti A, Griss J, Mayer G, Eisenacher M, et al. 2019. The PRIDE database and related tools and resources in 2019: improving support for quantification data. Nucleic Acids Res 47: D442–D450.

48. Reams A, Roth JR. 2015. Mechanisms of Gene Duplication and Amplification. Cold Spring Harb Perspect Biol 7: 1–25.

49. SaiSree L, Reddy M, Gowrishankar J. 2001. IS186 insertion at a hot spot in the lon promoter as a basis for Lon protease deficiency of Escherichia coli B: Identification of a consensus target sequence for IS186 transposition. J Bacteriol 183: 6943–6946.

50. Schedl P, Primakoff P. 1973. Mutants of Escherichia coli thermosensitive for the synthesis of transfer RNA. Proc Natl Acad Sci U S A 70: 2091–2095.

51. Sun L, Harris ME. 2007. Evidence that binding of C5 protein to P RNA enhances ribozyme catalysis by influencing active site metal ion affinity. RNA 13: 1505–1515.

52. Tallsjö A, Kirsebom LA. 1993. Product release is a rate-limiting step during cleavage by the catalytic RNA subunit of Escherichia coil RNase P. Nucleic Acids Res 21: 1686.

53. Thomason LC, Costantino N, Court DL. 2007. E. coli Genome Manipulation by P1 Transduction . Curr Protoc Mol Biol 79.

54. Tlsty TD, Albertini AM, Miller JH. 1984. Gene amplification in the lac region of E. coli. Cell 37: 217–224.

55. Tsilibaris V, Maenhaut-Michel G, Van Melderen L. 2006. Biological roles of the Lon ATP-dependent protease. Res Microbiol 157: 701–713.

56. Van Melderen L, Aertsen A. 2009. Regulation and quality control by Lon-dependent proteolysis. Res Microbiol 160: 645–651.

57. Vioque A, Arnez J, Altman S. 1988. Protein-RNA interactions in the RNase P holoenzyme from Escherichia coli. J Mol Biol 202: 835–848.

58. Wohlever ML, Baker TA, Sauer RT. 2013. A mutation in the N domain of Escherichia coli lon stabilizes dodecamers and selectively alters degradation of model substrates. J Bacteriol 195: 5622–5628.

